# Molecular insight toward efficient & robust design of vesiculated protein nano-cages

**DOI:** 10.1101/2025.03.15.643005

**Authors:** Shadi Rahnama, Mohammad Reza Ejtehadi

**Affiliations:** Sharif University of technology; Sharif University of Technology

## Abstract

Recently, recapitulation of viromimetic function in non-viral protein nanocages (PNCs) has emerged as a strategy to successfully encapsulate them in membrane vesicles. This method successfully evaded immune system detection. The mechanism responsible for triggering membrane budding and vesiculation remains elusive, primarily because the membrane initially interacts with a flat arrangement of proteins from nanocages (whether their shape is pyramidal, dodecahedron or icosahedron) and it is unclear how these seemingly flat protein arrangements can overcome the inherent mechanical resistance of the lipid bilayer to induce curvature. In this study, we considered a trimeric interface of a dodecahedron nanocage and explored the energetic and molecular role of its viromimetic module on protein nanocage packaging. Using a combination of all-atom and Martini coarse-graining molecular dynamics, we show that stronger highly basic region (HBR) promotes electrostatic sequestration of PIP2 lipids, known for their larger headgroups, around trimer binding sites, forming a PIP2 depletion zone in the central region of the trimer interface. Such lipid-sorting event resulted in membrane-thickness distribution with taller lipids accumulating toward the margins and shorter at the center of the trimer and inducing curvature to the lipid bilayer due to stretching and contraction events at different lipid interfaces. Our findings give molecular-level mechanistic insights into curvature generation and propagation in membranes induced by engineered PNC interactions, as well as a generic molecular design approach for clathrin-independent nanoparticle exocytosis.

## 1 Introduction

Protein nanocages hold immense promise for targeted drug delivery and immunologically stealthy therapeutics. A significant challenge hindering their clinical translation is the inefficient and uncontrolled encapsulation of these nanocages within membrane vesicles—vesiculation. To overcome this hurdle, researchers are increasingly turning to nature for inspiration, particularly to the elegant strategies employed by viruses for membrane manipulation and self-assembly. Years of research on viral replication and pathogenic pathways have yielded valuable insights into viral function[28]. Viruses employ an intrinsic mechanism to safeguard their genetic material, transport it to the extracellular space, evade immune responses upon entering target cells via interactions with receptors, and deliver it into the cell compartment. Through this process, they exploit self-assembled and distinctive protein structures known as capsids, which possess the capability to interact with host cells [13].

The capsid structure of viruses epitomizes the principles of protein self-assembly in nature. Through structural analysis, it’s revealed that this protein shell comprises a specific number of subunits that envelop and safeguard the viral genome [14]. Viral capsids possess inherent programming to target and penetrate host cells, evolved to facilitate nucleic acid exchange across different chemical environments [47].

Regarding their chemical and physical properties, capsids or natural nanocages exhibit stability against environmental pressures while remaining sensitive to signals or changes in the cellular environment, enabling the release of their cargo into the target microenvironment [33, 41]. The notable attributes of protein nanocages, including biocompatibility, functional diversity, biological manufacturing, and design flexibility through protein engineering, render them potent structures for various applications [49].

To date, viral capsids have found specialized applications. For instance, capsid bacteriophages are utilized in peptide display technology for synthesizing receptors for specific proteins, filamentous phages serve as templates for nanomaterial synthesis, and virus-like particles (VLPs) are employed in vaccine production, targeted drug delivery to cells, and the creation of bionanoreactors for material synthesis [49].

Despite the distinctive attributes of viruses, continuous endeavors persist to manipulate their protein sequences for the incorporation of novel or supplementary functionalities. Recent investigations have kindled optimism regarding the enhancement of nanocage architecture through the development of synthetic peptide sequences capable of self-assembling into structured protein nanocages [25, 26, 5, 18]. Analogous to their natural counterparts, peptide aggregation in these synthetic constructs arise from a diverse array of non-covalent interactions [52]. To enhance the utility of synthetic protein nanocages in diagnostic and therapeutic realms, as well as to modulate immune responses, they have been conjugated with diverse moieties—including sugars, lipids, nucleotides, and polymers—yielding hybrid protein scaffolds [7].

For instance, in recent research inspired by viral counterparts of synthetic protein nanocages, researchers introduced a modular method for conjugating structured nanocages with lipid membranes [50]. By emulating viruses that are encapsulated within lipid membranes, it is proposed that three fundamental capabilities exist in the genetic coding of any progenitor, conferring upon it the ability to be released into membrane vesicles, namely:

1. Possibility of binding to the membrane (To overcome the energetic barrier for curvature induction)
2. Self-structuring
3. Utilization of the ESCRT machinery to induce membrane curvature during the final stages of budding

In a test case, membrane binding capability is conferred by incorporating a sequence of six basic amino acids derived from the N-myristoylated Gag structure of the HIV. Furthermore, a polypeptide consisting of 52 residues from Gag p6 was utilized to introduce membrane curvature induction in the final step. The test case is vesiculated, and despite the innovative concept of the idea and evaluation with diverse membrane binding and late budding machinery from various viruses, it yielded mixed results. In certain instances, combinations of genetic codes led to the encapsulation of 15 or more nanocages within a single vesicle, while in others, fewer nanocages were encapsulated. However, consistent encapsulation of a specific number of nanocages within vesicles was not observed in any instance [50]. Precise control over nanocage encapsulation is imperative for therapeutic applications, underscoring the need to elucidate the underlying causes.

The proposed technique, using a natural coating for administered agents and relying on single-celled self-organization for production, promises a distinctive position among drug packaging tools. Understanding virus roles in Origin of Life scenarios [15, 16] and genetic mutation [23, 27, 12] further motivates biomimetic approaches. Achieving artificial structures mimicking their natural counterparts could lead to unprecedented scientific advancements and deeper natural understanding.

Many studies have explored the formation of inward tubular invaginations, predominantly focusing on proteins with homogeneous shapes characterized by high curvature, such as spheres, caps, cylinders, and ovals [39, 45, 37, 32, 4, 40, 53, 21, 29, 46, 54]. In contrast, nanocages often possess outer surfaces that are largely hollow, and their interacting surfaces with membranes are typically flat, unlike the aforementioned geometries. Furthermore, it should be noted that previous research did not always consider the atomic details of proteins. In this article, however, we emphasize the molecular structure of nanocages as a crucial element in understanding the mechanism of membrane curvature.

The nanocage in this study is a synthetic dodecahedron [18], with a consistently flat, hollow pentagonal interface (Figure 1). Each pentagonal structure comprises five trimers, with each trimer located at a vertex. Given that the bending rigidity of the nanocage is higher than that of the lipid bilayer [50, 18], understanding the specific effects of each trimer on the membrane is crucial for addressing their cooperation in inducing membrane curvature. Therefore, elucidating the underlying mechanism of invagination can be accomplished by investigating the induction mechanism of the system’s smallest building block: the trimers. These trimeric protein scaffolds possess a completely flat configuration; they are positioned at the corners of a pentagon (figure 1.a). In this study, we explore the significance of this spatial arrangement of trimers in the formation of vesiculated nanocages. To elucidate the curvature mechanism, we utilized coarse-grained and all-atom molecular dynamics simulations, which revealed that lipid sorting is the primary cause of membrane curvature by these flat structures. This process hints at the use of biophysical ‘cheap tricks’—an energy-efficient strategy that leverages the natural propensity of lipids to self-reorganize without substantial energy input [43, 31]. This sorting involves attracting longer lipids to the trimer corners and organizing shorter lipids in the interstitial spaces.

**Figure 1:**
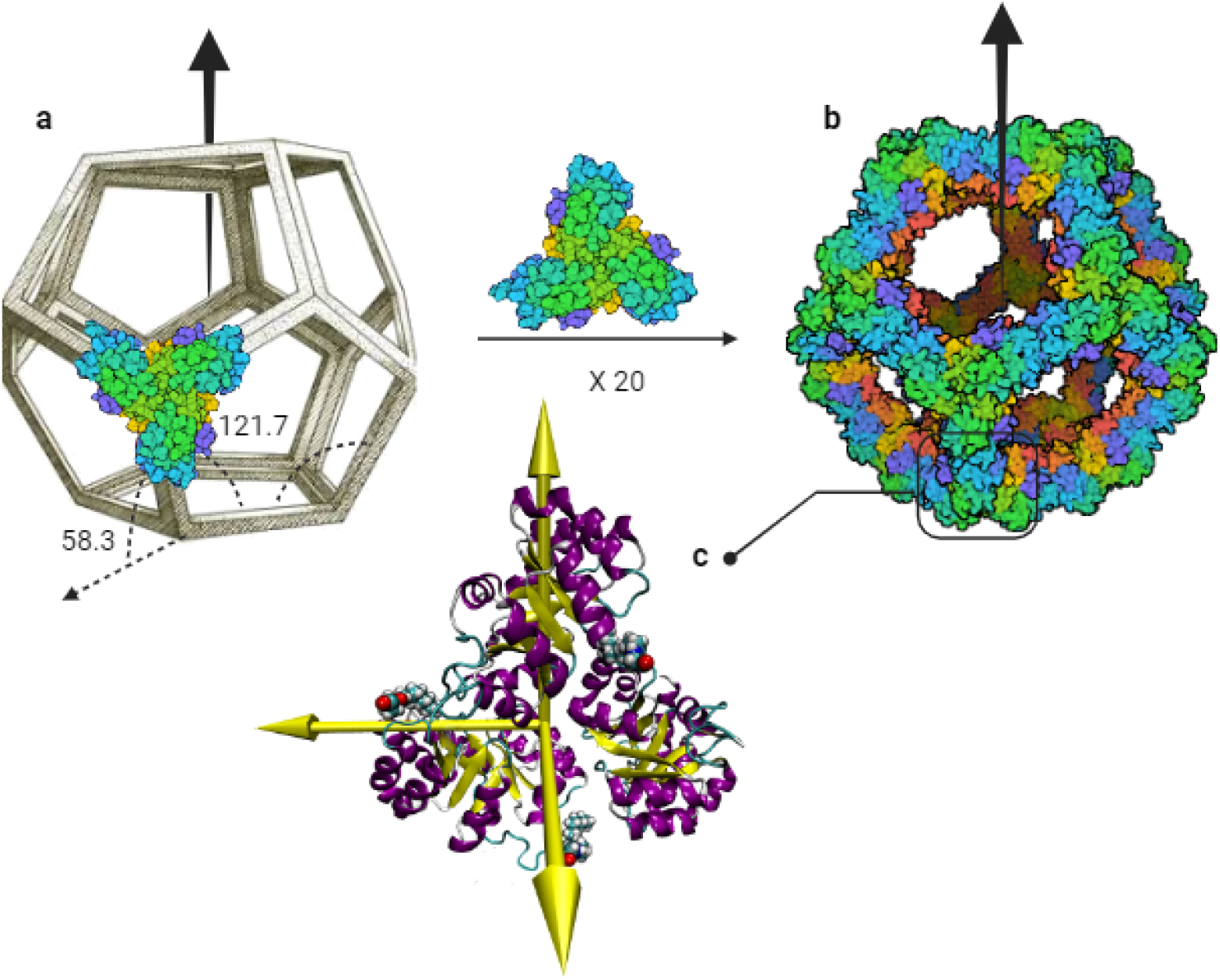
**A**. synthetic dodecahedron and an aligned trimeric building block, showing the angles of the landing surface in relation to the interface. **B**. A complete nanocage composing from 20 trimeric building blocks. **C**. Atomistic detail of a trimeric building block. Positioning at 58.3 degrees in relation to the interface places its lipid tail in a trigger position for anchoring to the lipid bilayer. The yellow axes are principle axes of the trimer.

## 2 Material and Methods

The initial coordinates of the protein nanocage were derived from PDB entry 5KP9. Unresolved amino acids, due to the low resolution (5Å) of electron microscopy, were simulated using Modeller [51] in Chimera [36] and incorporated into the original structure. The N-terminal region, which interacts with the membrane, includes a lipid tail. Five different approaches were used to modify the charge and lipidation state of the N-terminal region utilizing VMD [19] software and TCL [35] scripts. In Model I (figure 2), a Myristic acid tail was added to the N-terminal, following the model proposed by Vottler [50]; this serves as our native model, with the other models generated for comparison. In Model II (figure 2), the Myristic acid tail was replaced by a Palmitic acid lipid chain. Model III (figure 2) retained the myristoylation signal, as in Model I, but the first five amino acids of the HIV-1 Gag protein were replaced with a poly-positive lysine (K4) chain to introduce a strong positive charge. Models IV and V were designed to explore the role of an additional lipid tail at the N-terminal binding site. Model IV (figure 2) included an extra Myristic acid tail at Glycine14, compared to Model I, while Model V (figure 2) was identical to Model IV except that the 2-6 Gag sequence was replaced with a poly-neutralized Alanine sequence. The H++ algorithm [2] was used to estimate the pKa values of the ionizable amino acids of the protein at physiological pH. The calculated values indicated that all nine histidines present in the protein adopt a neutral protonation state. The lipid membrane was constructed using the Charm-GUI online platform [22]. The lipid composition was modeled to reflect the proportions observed in the HIV virus capsid, specifically 20% PIP2, 13% POPC, and 67% POPE [1]. The membrane was constructed with dimensions of 10 × 10 nanometers. The corresponding image, along with the structures of the constituent lipids, is presented in Figure 3.

**Figure 2:**
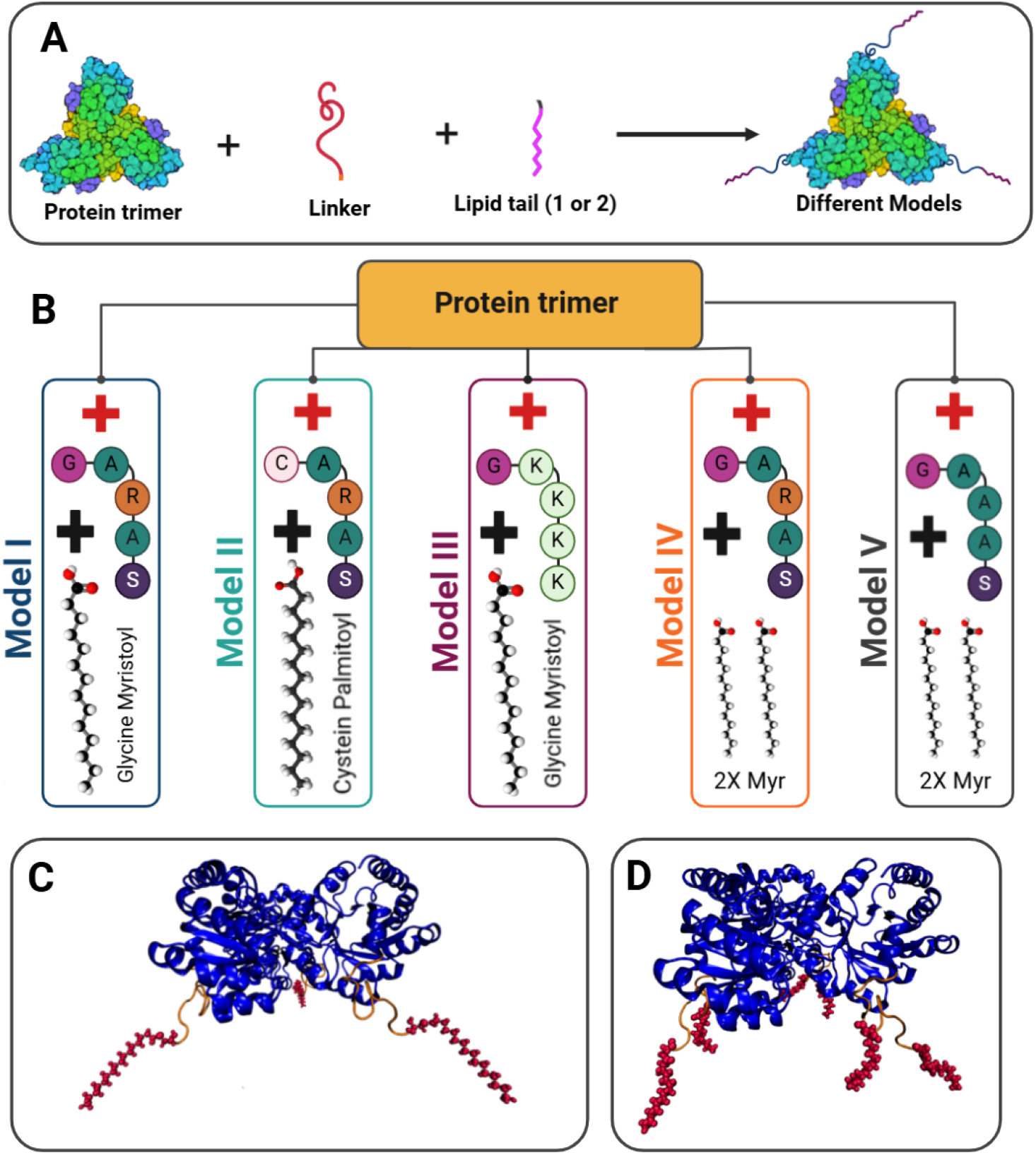
(A) Five different models of protein modification used in this study, created by adding a linker and one or two lipid tails to the primary protein trimer. (B) Atomistic differences between the models: **Model I** includes a single Myristic acid tail at the N-terminal and serves as the native model, with the linker sequence identical to the 2-6 *HIV-1 Gag* peptide sequence. **Model II** replaces the Myristic acid tail with a Palmitic acid chain. **Model III** retains the Myristic acid tail but introduces a strong positive charge by replacing the 2-6 Gag sequence with a poly-lysine (K4) chain. **Model IV** adds a second Myristic acid tail at Glycine14, while **Model V** is similar to Model IV but replaces the 2-6 Gag sequence with a poly-Alanine sequence, neutralizing the charge. (C) All-atom representation of Model III (The first three models are almost identical to this model in appearance, disregarding the details). (D) All-atom representation of Model IV (Model V is almost identical to this model in appearance, disregarding the details).

**Figure 3:**
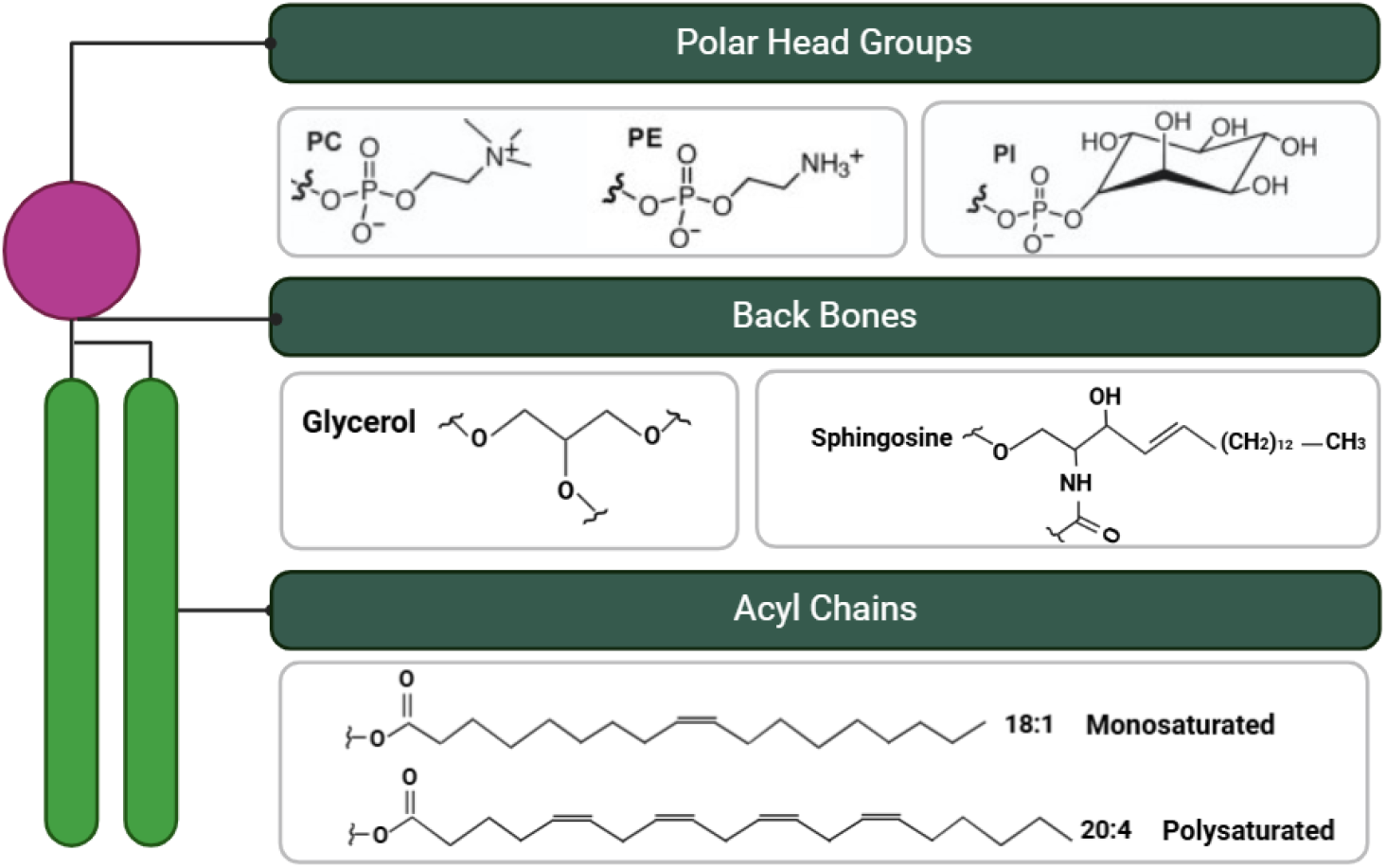
Structural representation of lipids used in this study. The lipids differ in their acyl chain length and degree of saturation: The monounsaturated lipids (POPC, POPE) contain 18-carbon acyl chains, while the polyunsaturated PiP2 lipids feature longer 20-carbon acyl chains. Also the larger polar head group is characteristic of PIP2 (Phosphatidylinositol 4,5-bisphosphate).

### 2.1 CG model

The membrane and protein trimers were mapped into a coarse-grained (CG) representation using the RBCG module in VMD. This module employs a coarse-graining force field based on the MARTINI force field [30], with modifications introduced by Shih and colleagues [44]. To preserve the secondary and tertiary structures of the proteins [3], an elastic network model was applied with a constant force of 0.597 kcal/(mol·Å^2^) between BAS beads separated by distances in the range of 4 Å to 10 Å, excluding the first 24 N-terminal residues (GLY-2 to LYS-25). Coarse-grained (CG) representations for the aliphatic moieties of the Myr and Pal groups were derived from established parameters within the MARTINI force field, including those used for lipids as recommended in the literature [9].

Water molecules, antifreeze beads, and counter-ions were added separately to each component of the system (membrane and protein trimers), which were equilibrated individually for 1 µs. Following equilibration, the systems were combined, with each protein model placed 12 Åaway from the surface of the membrane. Solvation and ionization were then performed for the combined systems resulting in simulation box with approximate size of 120 × 115 × 145 Åand containing approximately 15500 beads.

All coarse-grained simulations were carried out using the NAMD2.12 software [38]. Prior to the production simulations, the final structures of all models were minimized for 5000 steps to eliminate any steric clashes. Periodic boundary conditions were applied under the NPT ensemble, utilizing the Langevin thermostat and the Nosé-Hoover Langevin piston method to maintain a temperature of 310 K and a pressure of 1 bar. The Langevin thermostat damping coefficient was set to 5 ps, with a barostat time constant of 1000 fs and a collapse time constant of 500 fs. A time step of 10 fs was used throughout the simulations. Non-bonded interactions were calculated with a 1-2 exclusion and a cutoff distance of 12 Å.

For each model, two replicas were simulated, each undergoing unconstrained evolution for approximately 0.5 µs. The simulation results were analyzed using VMD software and custom Python scripts.

### 2.2 All-atom model

All-atom simulation just done for free energy calculation. All-atom setups for Model I were obtained by reverse mapping from its coarse-grained MARTINI model, followed by an annealing process to elevate the temperature to physiological levels. The resulting structures were used as the initial configuration for all-atom modeling, with ten replicas designed for free energy calculations. Each system was solvated using the TIP3P water model and neutralized to 150 mM NaCl, resulting in a water box of approximately 132 × 132 × 158 Å^3^, containing around 192,000 atoms. CHARMM36m force field parameters [48] were applied, and molecular dynamics simulations were conducted using NAMD 2.12 [38]. Periodic boundary conditions under the NPT ensemble maintained the temperature and pressure at 310 K and 1 bar using the Langevin thermostat and Nosé-Hoover Langevin piston. A 12 Å cutoff was used for short-range van der Waals interactions, and long-range electrostatics were calculated via the Particle Mesh Ewald method. The R-RESPA multiple time step scheme, with a 2 fs time step, was used for integrating the equations of motion. Structures were minimized for 5,000 steps to remove steric clashes, followed by equilibration: first for 0.5 ns in the NVT ensemble with restrained lipid tails, then for another 0.5 ns in the NPT ensemble under the same restraints. Given prior coarse-grained equilibrium, only an additional 1 ns of unrestrained all-atom modeling was necessary before performing free energy calculations.

### 2.3 Free Energy Calculation

The anchoring process of the trimer is typically accompanied by significant conformational changes in the protein. The total calculated potential of mean force (PMF) throughout the evolution of this process is composed of both the binding free energy of the protein to the lipid membrane and its associated conformational changes. To isolate the binding free energy component, we utilized the Collective Variables (colvars) [6] module in NAMD, which effectively reduces the protein’s high-dimensional degrees of freedom into a manageable set of key parameters. By defining biasing potentials, we carefully modulated the system’s dynamics in a controlled manner. In this study, we employed Steered Molecular Dynamics (SMD) to introduce biasing potentials, thereby enhancing sampling efficiency and facilitating exploration of the phase space along the reaction coordinate (Z). The protein trimer was pulled from the lipid membrane at a constant velocity of 0.0002 Å/timestep, with a virtual string attached to the protein’s center of mass, and a force constant of 7.5 *KCal/*(*mol* · Å^2^) was applied. Simultaneously, a counteracting restraint (*K* = 20 *KCal/*(*mol* · Å^2^)) was applied to the phosphate groups of the lipid molecules to prevent membrane translocation in response to the biasing potential. The work performed during this non-equilibrium process is related to the equilibrium PMF via Jarzynski’s equality [20], given by:

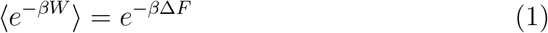

where 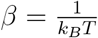, *W* is the work done on the system, and Δ*F* is the change in free energy.

## 3 Results and Discussion

Regardless of the membrane-binding motif, the majority of the replica simulations reach an equilibrium state after 200 ns of simulation time, as indicated by the mean RMSD of each model (Figure Appendix 18). In nearly all models, the mean RMSD stabilizes, reaching a plateau around 150 ns of simulation. Therefore, subsequent analyses were conducted post-equilibration.

To assess the impact of the motifs on each model’s interaction with the membrane, we first evaluated the number of acyl tails that penetrated the membrane (Figure 4). Model III exhibited the highest efficiency, with all three acyl tails inserting into the membrane on average across both repetitions. In this model, the poly-positive lysine (*K*_4_) chain of the linker introduced the strongest charge in the membrane-binding motifs among the models, resulting in the strongest electrostatic interaction with the membrane. For Models I and II, which incorporate the HIV-derived electrostatic motif, the average numbers of inserted tails were 2.49 ±0.02 and 2.35 ±0.01, respectively (Figure 4).

**Figure 4:**
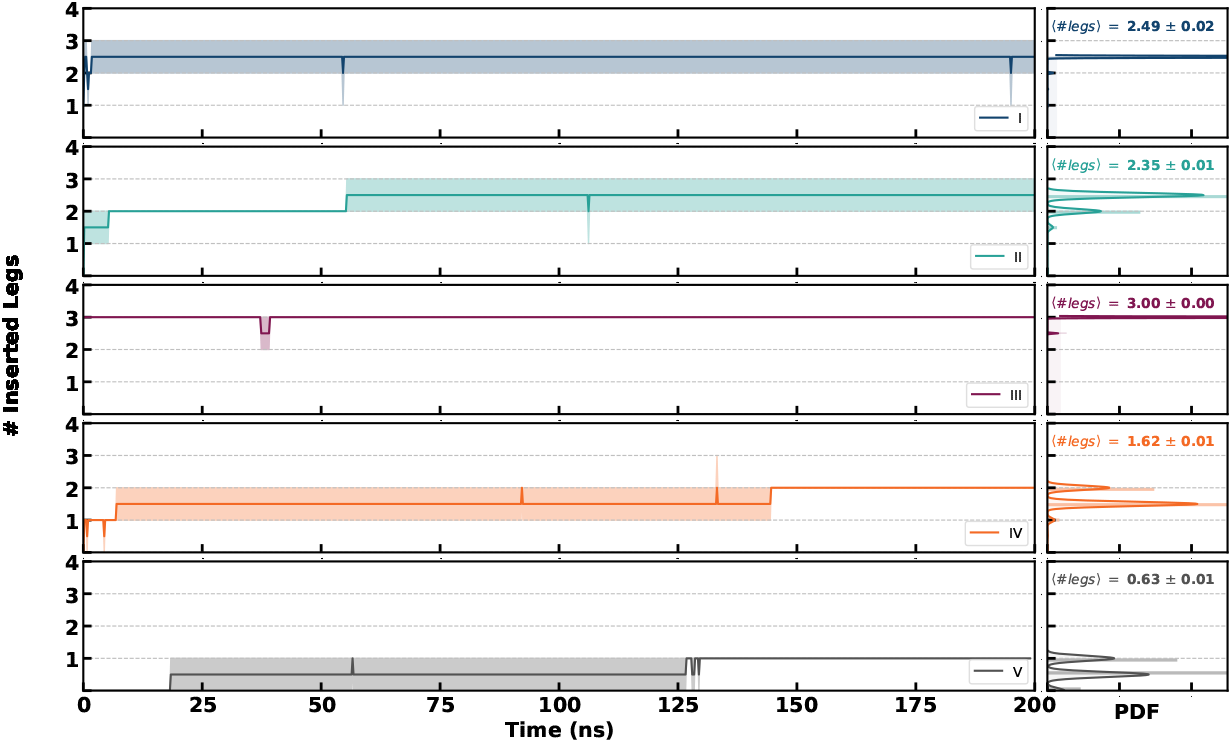
Number of monomers in each model that successfully inserted at least one tail into the membrane over the simulation time. The left panels display the number of inserted tails (legs) as a function of time, while the right panels show the corresponding probability density functions (PDF) of the inserted tails for each model (I to V). The average number of inserted tails (⟨#legs⟩) is indicated for each model, highlighting variations in tail insertion dynamics across different systems.

In the remaining models, which have two acyl tails per monomer, the results differ. In Model IV, the tail attached to the first residue has an electrostatic motif similar to Models I and II, while the tail on the eleventh residue does not. This model exhibited less success in membrane interaction than Models I and II. The tail on the eleventh residue showed significant fluctuations, interfering with the first tail and often leading to both being drawn into the hydrophobic cavity of the protein. As a result, an average of 1.62 ± 0.01 tail penetrations into the membrane was recorded in this model (Figure 4).

The situation worsened in Model V, where the charged motif was replaced with a neutral series of Alanines. Similar disruptions to those observed in Model IV were also present, leading to an even lower average lipid tail insertion, dropping below one to 0.63 ± 0.01 (Figure 4).

The overall results clearly indicate that the electrostatic charge of the membranebinding motif at the N-terminus of the monomers significantly enhances the tail insertion phenomenon into the membrane.

Curvature induction analysis across different models reveals a spectrum of curvature-inducing capabilities, as quantified in Figure 5. To visually exemplify this curvature induction, Figure 6 illustrates the final snapshot of the system for Model III, which exhibited the most pronounced curvature in our analysis. This snapshot reveals that the protein trimer in Model III indeed induces a noticeable local curvature on the bilayer surface. To provide a clearer depiction of this induced curvature, Figure 6B presents the membrane surface fitted to a fifth-degree polynomial, as detailed in the appendix.

**Figure 5:**
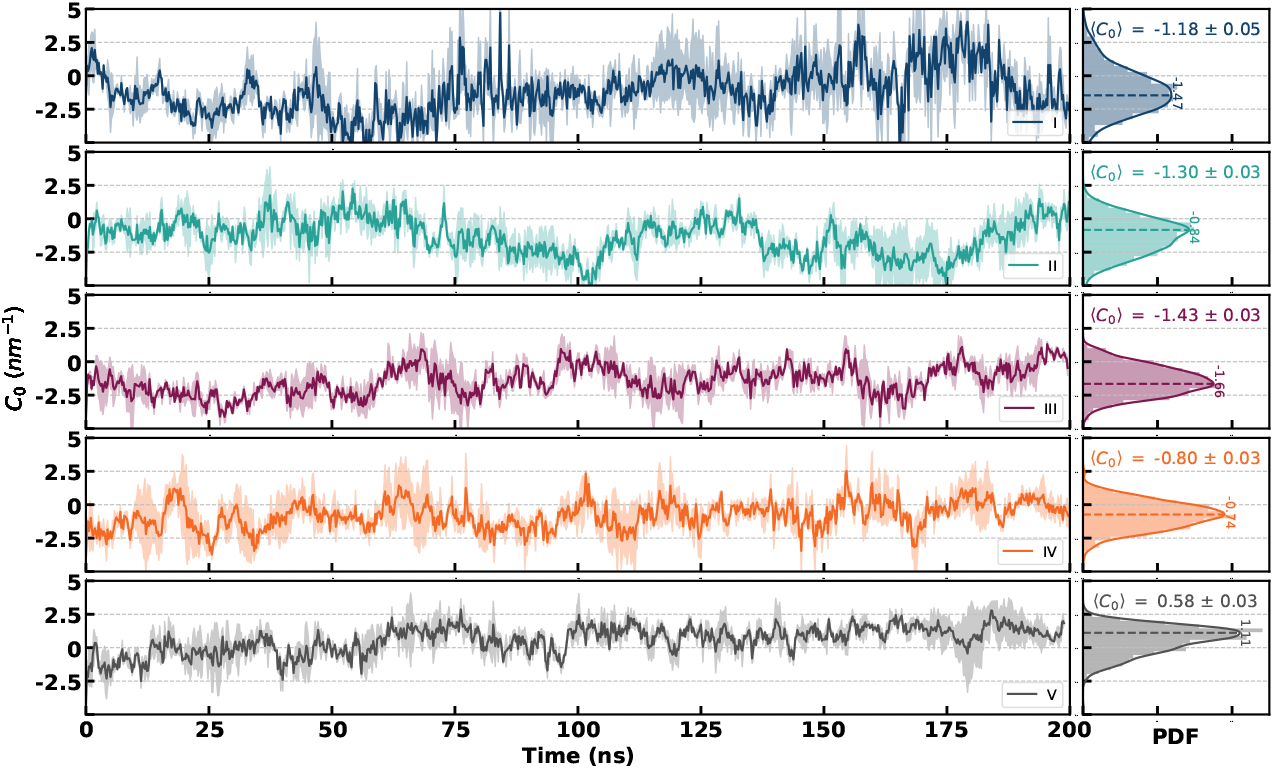
Plot of local membrane curvature variations (*C*_0_) over time in different protein models. Each subplot shows the time evolution of *C*_0_ (left) alongside the corresponding probability density function (PDF) (right). The average curvature (⟨*C*_0_⟩) is indicated for each model, revealing distinct curvature dynamics across the five protein models (I to V), with both positive and negative curvature trends observed.

**Figure 6:**
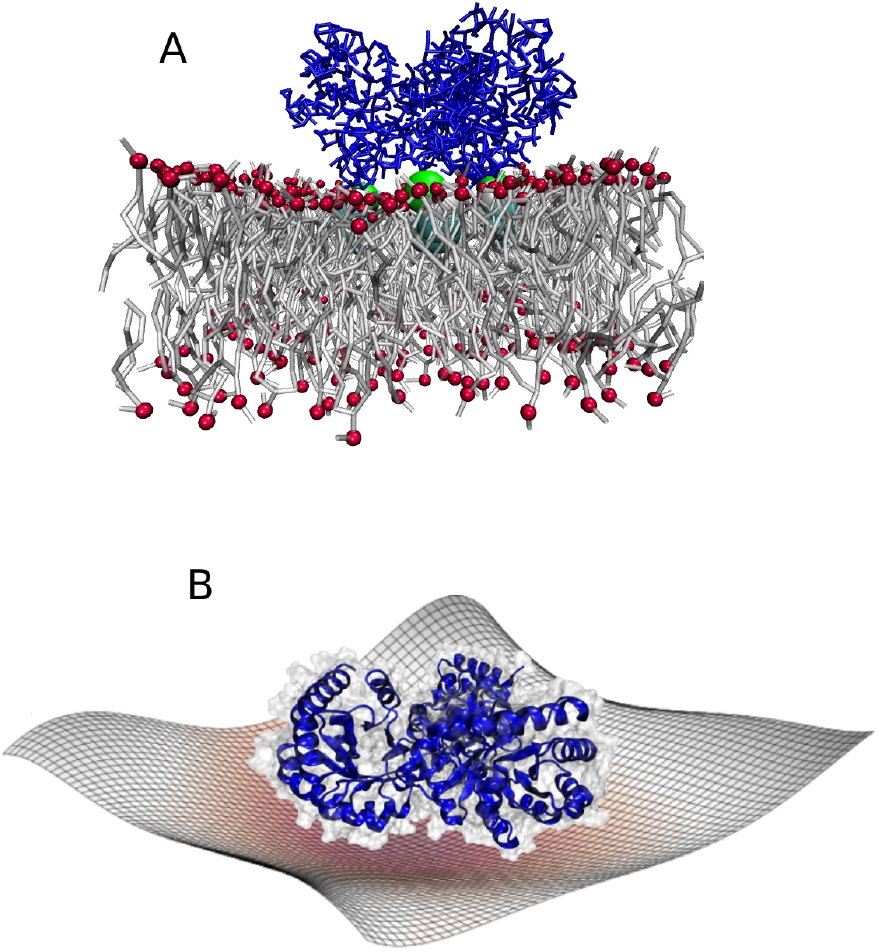
(A) Membrane-protein complex inducing curvature in the membrane. (B) Visualization of the membrane surface showing the induced curvature more clearly, with the surface shape highlighted using polynomial fitting.

Quantitative analysis of curvature induction (Figure 5) illustrates that Model III induces the most significant curvature in the membrane (⟨*C*_0_⟩ = −1.43± 0.03 nm^−1^), correlating with the highest number of monomers successfully inserting their lipid tails into the membrane. This suggests that efficient penetration of monomers is a key driver of membrane deformation. In contrast, Model V results in the lowest curvature (⟨*C*_0_⟩ = 0.58 ± 0.03nm^−1^), indicating a weaker influence on the membrane structure, likely due to its minimal tail insertion (Figure 4). This observation underscores the importance of monomer insertion in determining the extent of induced curvature.

Models I and II induce intermediate levels of membrane curvature, with ⟨*C*_0_⟩ = −1.18 ±0.05, nm^−1^ and ⟨*C*_0_⟩ = −1.30 ± 0.03nm^−1^, respectively. Interestingly, although Model I shows a higher average number of tail insertions into the membrane (2.49 ±0.02 insertions) compared to Model II (2.35 ±0.01 insertions), it results in a lower curvature. This apparent contradiction indicates that membrane curvature is influenced by factors beyond just the number of tail insertions. The specific nature of the membrane-protein interactions, as well as the distribution and physical properties of the inserted tails, likely play a significant role in modulating the overall curvature, which will be discussed in more depth later on.

Model IV shows a relatively lower curvature (⟨*C*_0_⟩ = −0.8 ± 0.03nm^−1^), aligning with its reduced capacity for tail insertion. This model, along with Model V, demonstrates that insufficient penetration of monomers into the membrane correlates with a decrease in membrane deformation.

Overall, our results show that the degree of induced curvature is thus directly linked to the inserted tail stoichiometry. But the mechanism that how tail insertion to the membrane could induce curvature to it is still unclear. To gain a deeper understanding of underlaying molecular mechanism of curvature induction, we next investigated the correlative behaviour of membrane interacting motifs of protein and lipids of membrane.

To achieve this, we first calculated the distribution function of the presence of protein’s “membrane-binding motifs” and PIP2 lipids in the XY plane (Figure 7). This allows us to identify potential spatial correlations that could provide insights into the mechanisms of curvature induction. In Figure 7, the probability distribution of the presence of PIP2 molecules is presented in the left panels, while for protein’s binding motifs in the right ones. Preliminary analysis of Figure 7 reveals a distinct spatial correlation between these two quantities. For instance, in the model III, three prominent orange-reddish regions are visible at the centers (0, 20), (0, − 20), and (0, 40) in the right panel, representing highly probable regions for protein motif insertion. These regions coincide with the high-probability regions of the presence of PIP2 in the left panel. Seemingly, these hotspot regions of protein motifs act as absorption points for PIP2 lipids, leading to areas with a high probability of PIP2 presence (PIP2-enriched regions) in the left panel. Conversely, colder-colored regions in the left panel indicate areas that appear to be depleted of PIP2 lipids. This pattern is observed to varying degrees across other models as well. However, it remains unclear whether this relationship between protein and membrane components is consistent across different systems.

**Figure 7:**
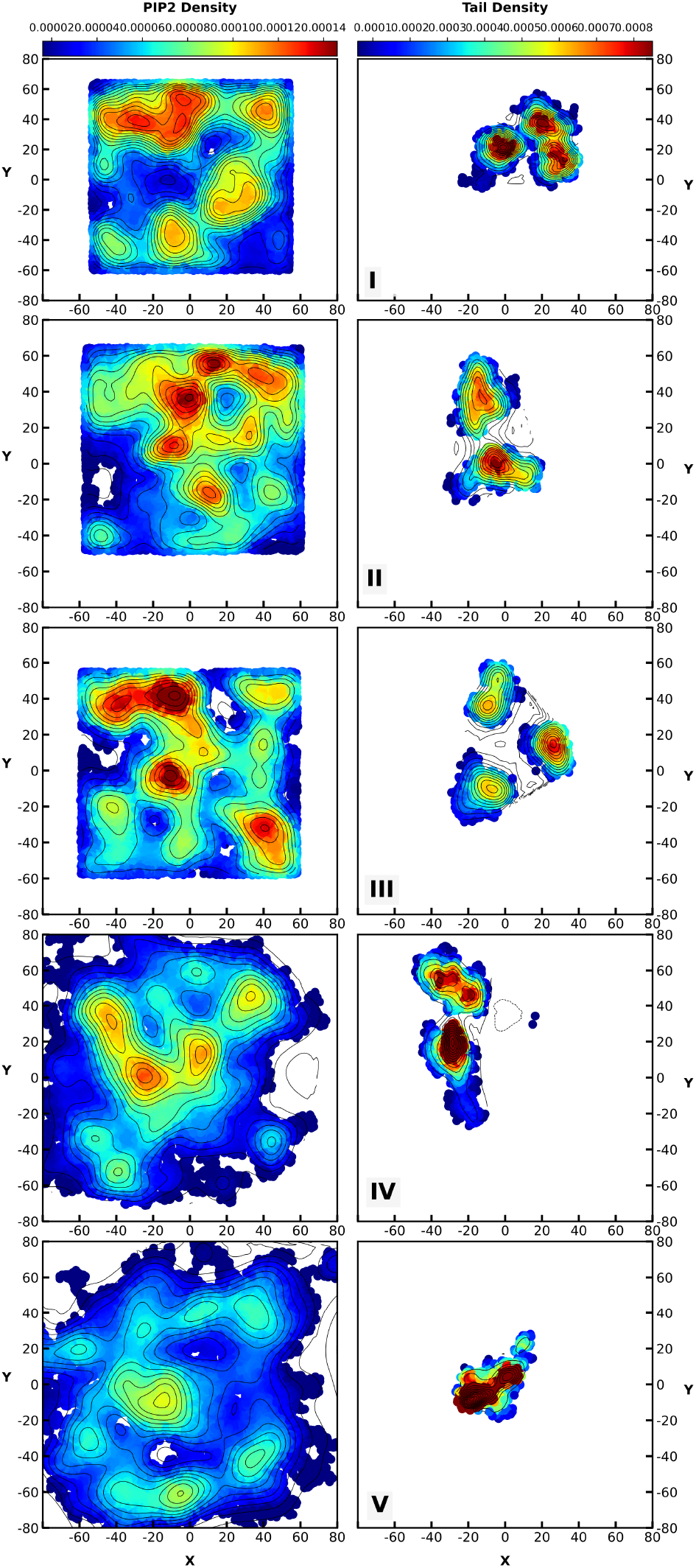
Probability distribution of PIP2 molecules (left) compared to the probability distribution of protein tail insertion (right) in the membrane surface for different models. The color scale represents the probability density, with red regions indicating higher density and blue regions indicating lower density. The spatial correlations between PIP2 lipid presence and tail insertion points suggest a significant relationship across different models.

To address this, we defined a metric called the “relative probability” (*f*_*τϕ*_) to quantify the overlap between the distributions of the motifs and PIP2 lipids on the membrane surface:

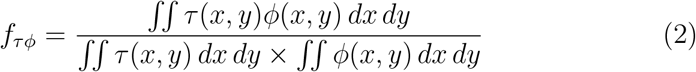

Here, *τ* (*x, y*) represents the probability distribution of the motifs, and *ϕ*(*x, y*) represents the probability distribution of PIP2 lipids. This formula measures the degree of co-localization between the motifs and PIP2 lipids. A higher value of *f*_*τϕ*_ indicates a stronger spatial correlation, implying more effective formation of absorbent points, while a lower value indicates vice versa.

To quantify the correlation between the induced curvature (*C*_0_) and the relative probability (*f*_*τϕ*_), we plotted the scatter plot of these variables in Figure 8. The horizontal axis represents the induced curvature, while the vertical axis shows *f*_*τϕ*_. Each point on the scatter plot corresponds to a specific model, indicating the values of both variables. The data suggest that as the induced curvature becomes more negative, the value of *f*_*τϕ*_ increases.

**Figure 8:**
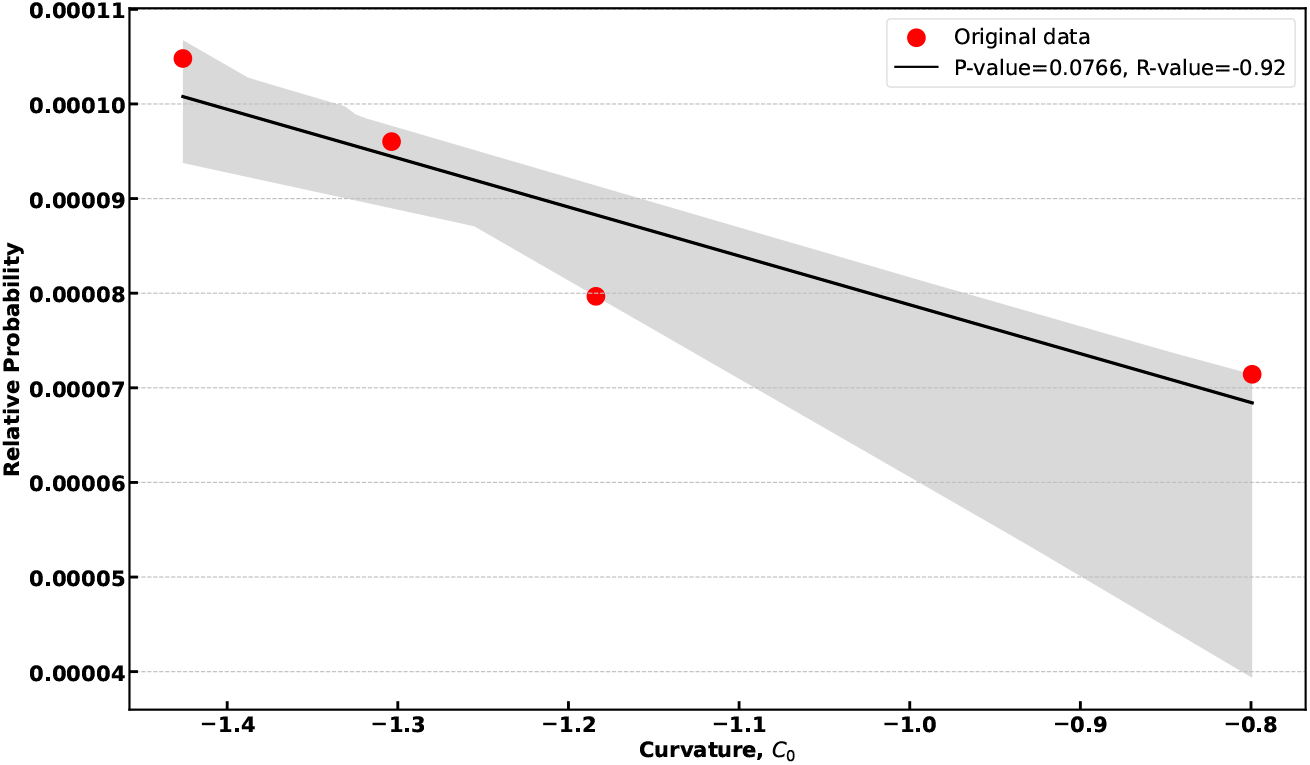
Correlation between induced curvature (*C*_0_) and relative probability (*f*_*τϕ*_). The data shows an inverse relationship (R-value = -0.92, P-value = 0.0766), with the shaded area representing the 95% confidence interval. The red data points correspond to original values from different models, and the black line represents the linear regression fit.

Performing linear regression on the data yielded a correlation coefficient of corr(*C*_0_, *f*_*τϕ*_) = − 0.92 and a p-value of 0.07 (Figure 8), further confirming the strong inverse relationship between these variables and the consistent correlation between the presence probability of protein motifs and PIP2 lipids across different models. Model V was omitted from this correlation analysis due to its inability to insert more than one lipid tail into the membrane.The reasons for this exclusion are discussed subsequently, following the detailed explanation of the curvature induction mechanism. This ordering ensures that the explanation for Model V’s distinct behavior is presented with the necessary mechanistic context.

A closer examination of the molecular structure of the PIP2-enriched and PIP2-depleted regions brings us closer to identifying the underlying reason for the formation of this correlation: As depicted in Figure 3, PIP2 lipids feature multi-unsturated acyl chains composed of 20 carbon atoms, whereas POPC and POPE have shorter chains with 18 carbon atoms, with POPC being saturated and POPE monounsaturated. In typical phospholipids like POPC, the glycerol backbone is oriented approximately perpendicular to the membrane plane, while the head group lies parallel to it [17]. By contrast, the PIP2 head group aligns perpendicularly with respect to the membrane [8, 55]. The notable size and potentially upright structure of the PIP2 head group suggest that it may protrude more prominently into the aqueous phase compared to other phospholipids. These properties collectively make PIP2 “taller” than other membrane lipids.

Electrostatic sequestration between PIP2 lipids and positively charged motif of the protein induces phase separation within the membrane. This phenomenon exemplifies another of nature’s biophysical ‘cheap tricks’ [31], facilitating complex molecular rearrangements with minimal energy expenditure. Consequently, longer PIP2 lipids accumulate at the protein’s corners (corresponding to the vertices of an equilateral triangle where absorbent points or charged membrane-binding motifs are located), while the central region of the protein trimer becomes depleted of PIP2 (Figure 9). This region, in turn, becomes enriched with shorter lipids such as POPC and POPE, forming distinct domains. Such differential lipid distribution could be a key driver of membrane curvature. This phenomenon can be visually observed in Figure 9, which presents a topographic view of the membrane, color-coded to show PIP2 lipids (magenta) and protein myristoylate tails (green). As observed in Figure 9, regions of elevated membrane level are shown in purple, and these regions are also enriched in PIP2 lipids. Conversely, regions of lower membrane level are shown in white and exhibit a lower concentration of PIP2. Notably, the presence of the protein complex leads to a depletion of PIP2 in the membrane region interacting with the protein’s central area, and a corresponding enrichment of PIP2 in the peripheral regions surrounding the protein; this is clearly visualized in the figure.

**Figure 9:**
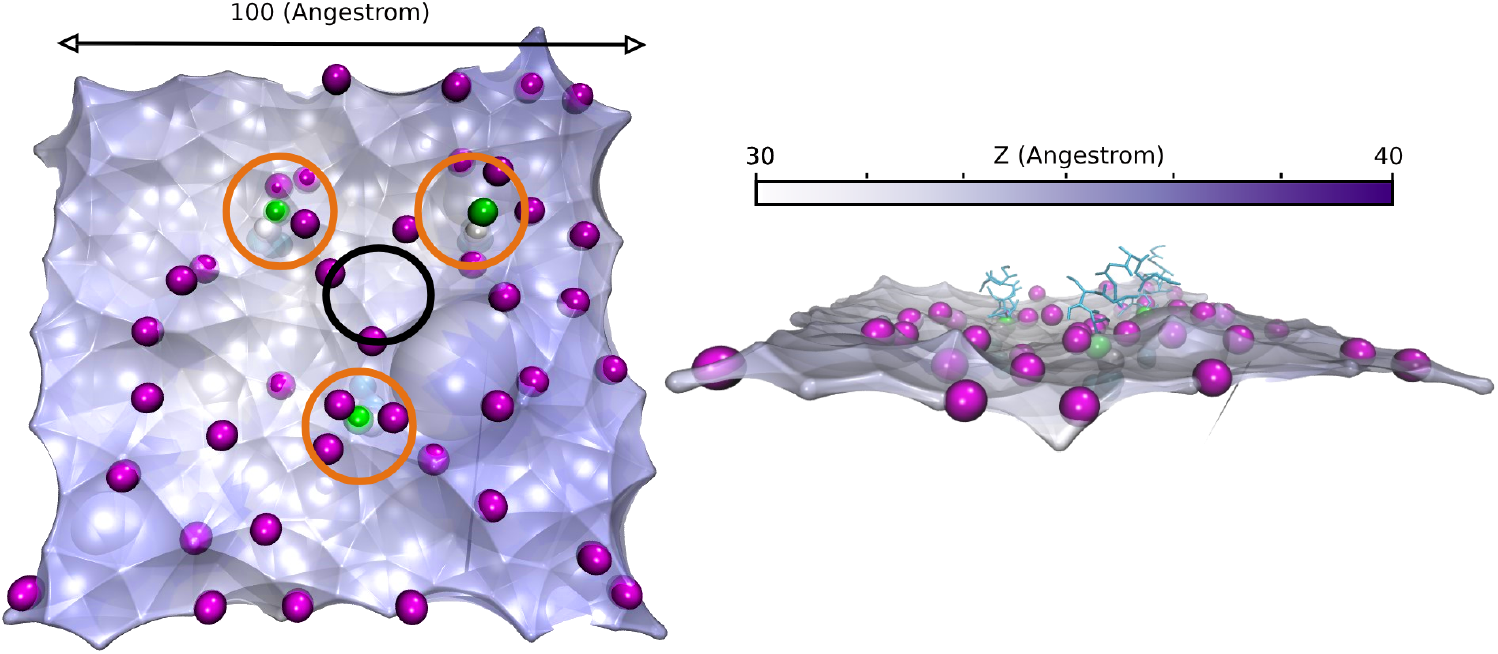
Visualization of membrane topography, showing PIP2 lipids (magenta) and myristate tails (green). Purple areas indicate elevated membrane levels (greater Z-depth) that correlate with higher PIP2 concentration, while white areas indicate lower membrane levels. This visualization demonstrates membrane deformation induced by PIP2 clustering and lipid sorting. The absorbing points were shown by orang circle and central region with black one.

The resulting membrane domains differ in molecular properties, including the thickness of the hydrophobic region. The height difference between these lipid domains resembles a step or bridge (Figure 11.A). This step exposes the hydrophobic parts of the longer lipids to the aqueous environment, which is energetically unfavorable. To mitigate this exposure, boundary lipids adjust by altering their orientations and lengths, effectively stretching or shrinking to cover the exposed hydrophobic regions, as shown in Figure 11.B [42, 24]. In this figure, two domains interact: one with taller lipids (black heads) and one with shorter lipids (white heads). In Panel A, the abrupt boundary leads to potential hydrophobic exposure. In Panel B, the shorter lipids stretch and the longer lipids compress, creating a gradual slope between the domains. This adaptation prevents direct hydrophobic exposure, maintaining membrane integrity. In a three-dimensional context, if this smooth transition occurs radially around a central point, it can result in an inward-curving cap-like structure. The taller lipids taper to align with the shorter lipids, forming a slope that converges inward, creating a cone-shaped structure with inward curvature.

Building upon our observation of PIP_2_ enrichment, this lipid sorting process appears to create regions of differential membrane thickness – with PIP_2_-rich areas potentially exhibiting increased thickness due to the larger PIP_2_ headgroup and chain properties, and PIP_2_-depleted regions thinning accordingly. To quantify this ‘relative thickness,’ which we hypothesize is mechanistically coupled to curvature induction, we calculated the metric *f*_*ϕt*_, defined as:

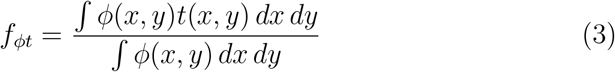

The relationship between induced curvature (*C*_0_) and relative thickness (*f*_*ϕt*_) across Models I-IV is presented in Figure 10. Although a linear fit reveals a weak correlation (*R*^2^ = 0.898, *p* = 0.102), the logistic fit provides a significantly better fit, as indicated by a substantially higher *R*^2^ value of 0.987 and a significantly lower *p*-value of 0.013. This nonlinear relationship suggests a threshold effect: at lower curvature values, relative thickness increases gradually, but beyond a critical curvature value, even small increments in curvature lead to a sharp rise in relative thickness. The decreasing trend in Figure 10 strongly supports the role of PIP_2_-driven lipid sorting and membrane thickness variations in curvature induction by protein trimers.

**Figure 10:**
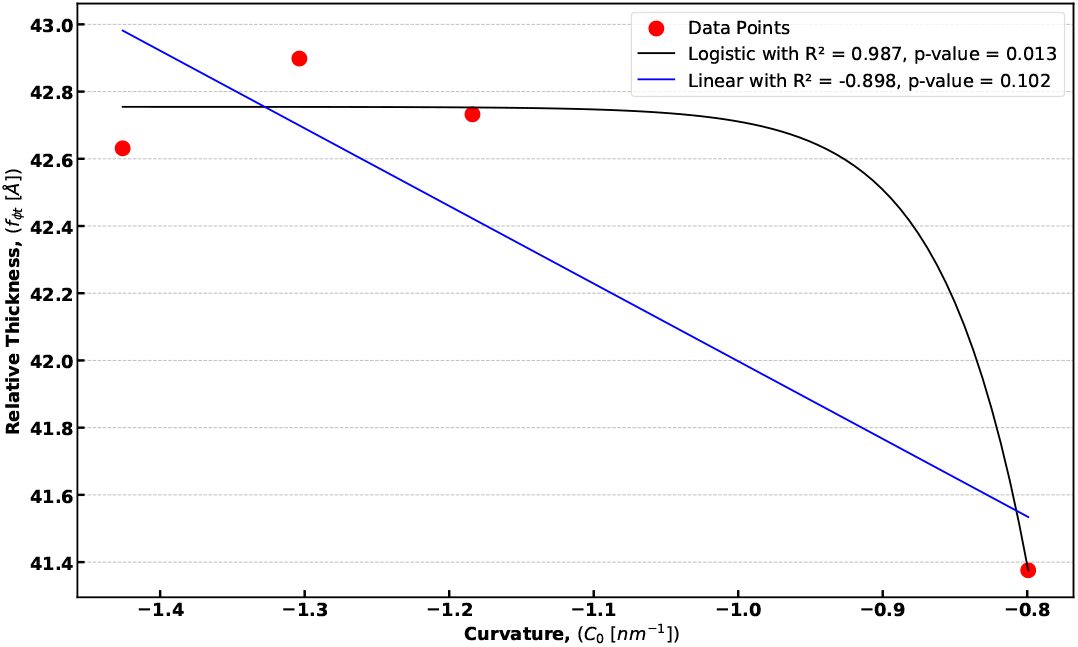
Relationship between Induced Membrane Curvature and Relative Membrane Thickness across Protein Models (I-V). Data points (red circles) represent measurements for Models I-V. The black dashed line shows the Logistic Regression Fit to the data (R^2^ = 0.987, p-value ≈ 0.013). For comparison, the blue line shows the Linear Regression Fit (R^2^ = 0.898, p-value ≈ 0.102). Curvature (*C*_0_) is given in units of nm^−1^, and Relative Thickness is presented with units of Å. The logistic model demonstrates a significantly better fit to the data (higher R^2^ and lower p-value) compared to the linear model, indicating a nonlinear relationship between induced membrane curvature and relative membrane thickness.

**Figure 11:**
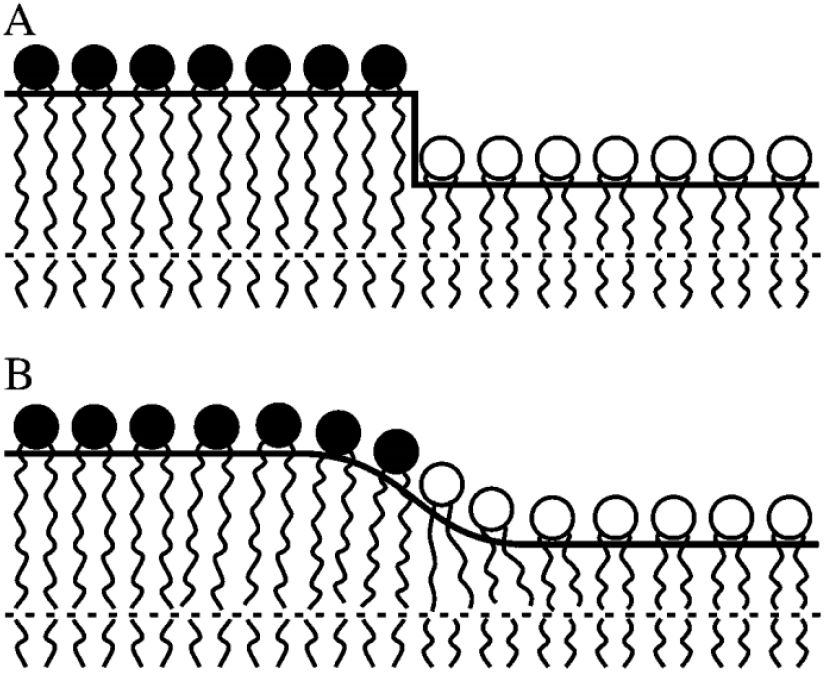
(A) A lipid step leading to hydrophobic exposure at the boundary between taller and shorter lipids. (B) Adjustment of boundary lipids to cover the hydrophobic region, with the longer lipids compressing and shorter lipids stretching to form a gradual slope, mitigating hydrophobic exposure [24].

In the explanation above, it was assumed that three tails could successfully insert into the membrane, leading to a cap-like shape (Figure 12.A). However, as previously shown, except for Model III, no other model could achieve this level of tail insertion. It is clear that when more than one leg inserts into the membrane (as in Models I, II, and IV), the membrane reshapes according to the described mechanism, resulting in a concave form or curvature induction (Figure 12.B).

**Figure 12:**
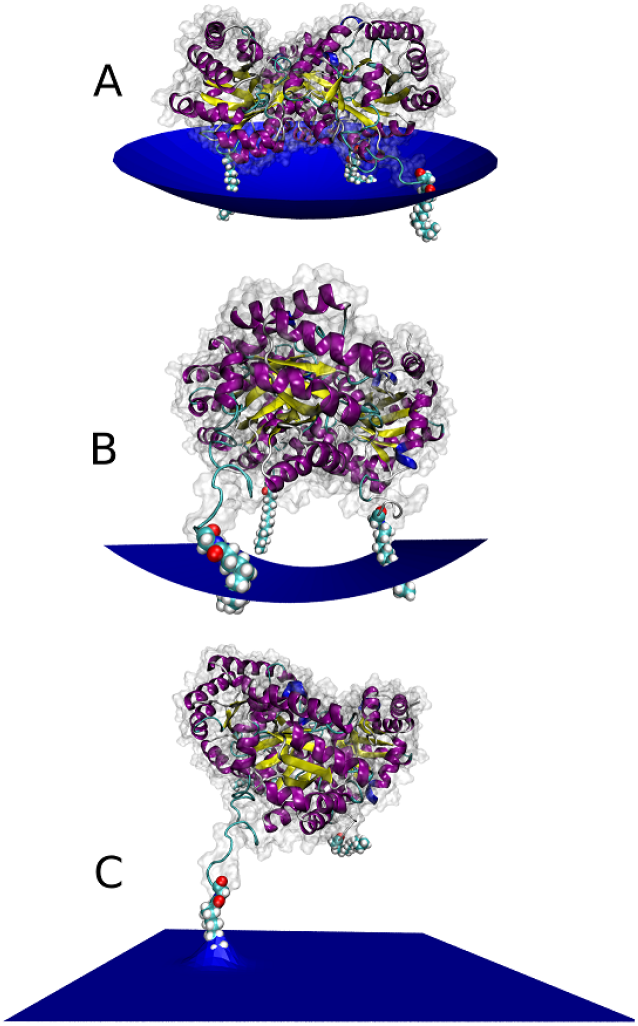
The model illustrates the effect of protein tail stoichiometry on membrane curvature. (A) Insertion of multiple tails results in a relatively flat or slightly curved membrane. (B) Insertion of two tails induces a prominent concave membrane curvature. (C) Insertion of a single tail causes localized upward deformation at the insertion point, with minimal overall curvature.

**Figure 13:**
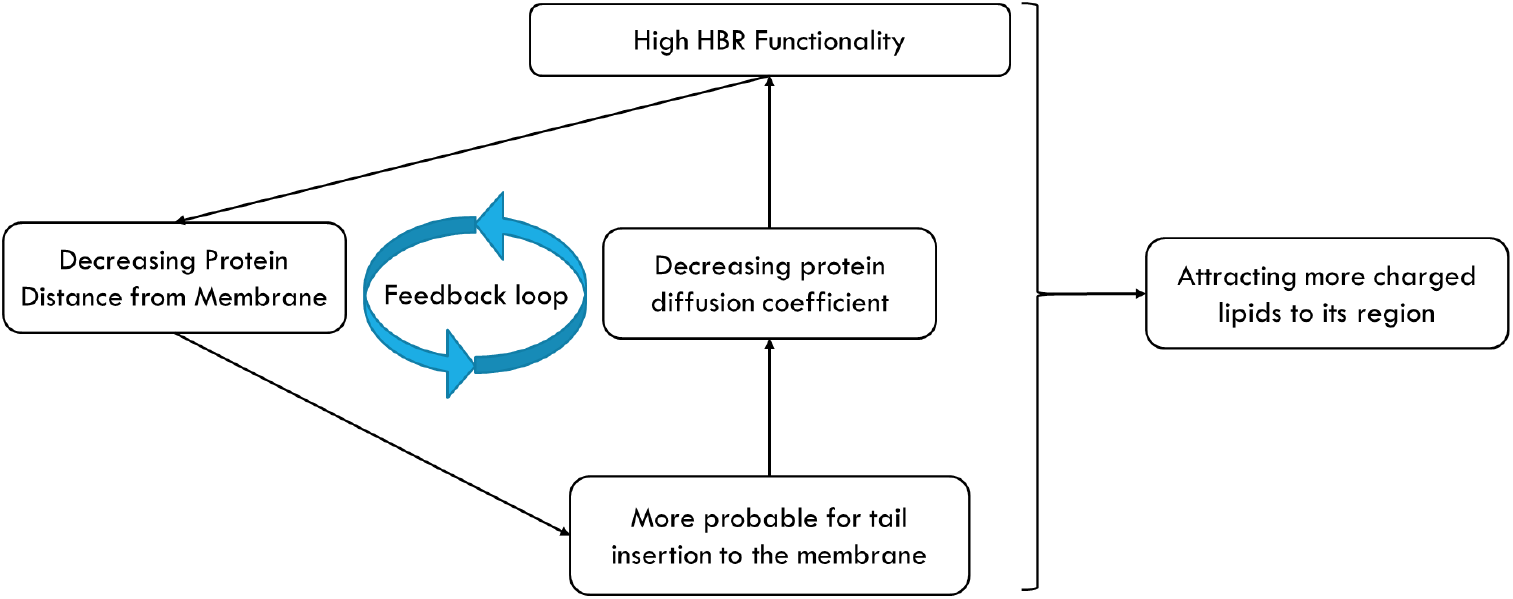
Diagram illustrating the synergistic feedback loop driven by a high HBR (Highly Basic Region) and the lipid tails of trimers in inducing curvature in the lipid membrane. Strong electrostatic interactions at the highly basic region promote lipid tail insertion into the membrane, which reduces protein diffusion and mobility. This restricted mobility facilitates the accumulation of negatively charged lipids around the protein, further decreasing the distance between the protein and the membrane and reinforcing the protein-lipid complex. This mutual reinforcement amplifies proteinmembrane interactions, leading to increased membrane curvature.

The situation is notably different when only one leg is successfully inserted into the membrane, as seen in some replicas of Model V. In this case, an uphill-like point forms near the leg insertion site, as only one absorption point is created, which is insufficient to produce a concave shape. For concavity to occur, at least two legs need to be inserted into the membrane.

It is also important to note that in Model V, no electrostatic interaction is present at the N-terminus of the protein trimers. According to the literature, proteins can successfully insert their lipid tails into the membrane when a large enough lipid packing defect is available [34]. The accumulation of PIP2 in certain regions may create these defects, allowing Model V to insert its tail into the membrane.

As discussed earlier, lipid reorganization mediated by electrostatic interactions between proteins (via the highly basic region, or HBR) and charged lipids is a key mechanism for inducing membrane curvature. However, proteins that do not incorporate their lipid tails into the membrane are less effective in initiating this process. The underlying reasons for this reduced efficacy are explored in the following sections.

Trajectory comparisons across various systems suggest that lipid tail insertion into the membrane initially reduces the lateral mobility of proteins on the membrane surface (Figure 14). Models with fewer lipid tails inserted (IV and V) exhibit greater protein mobility, reflected by higher diffusion coefficients (*D*_*IV*_ = 3.55 ×10^−8^cm^2^*/*S, *D*_*V*_ = 5.516 ×10^−8^cm^2^*/*S), while models I, II, and III show more restricted motion within the local lipid environment, corresponding to lower diffusion coefficients (*D*_*I*_ = 2.919 × 10^−8^cm^2^*/*S *> D*_*II*_ = 2.324 × 10^−8^cm^2^*/*S *> D*_*III*_ = 1.69816 × 10^−8^cm^2^*/*S).

**Figure 14:**
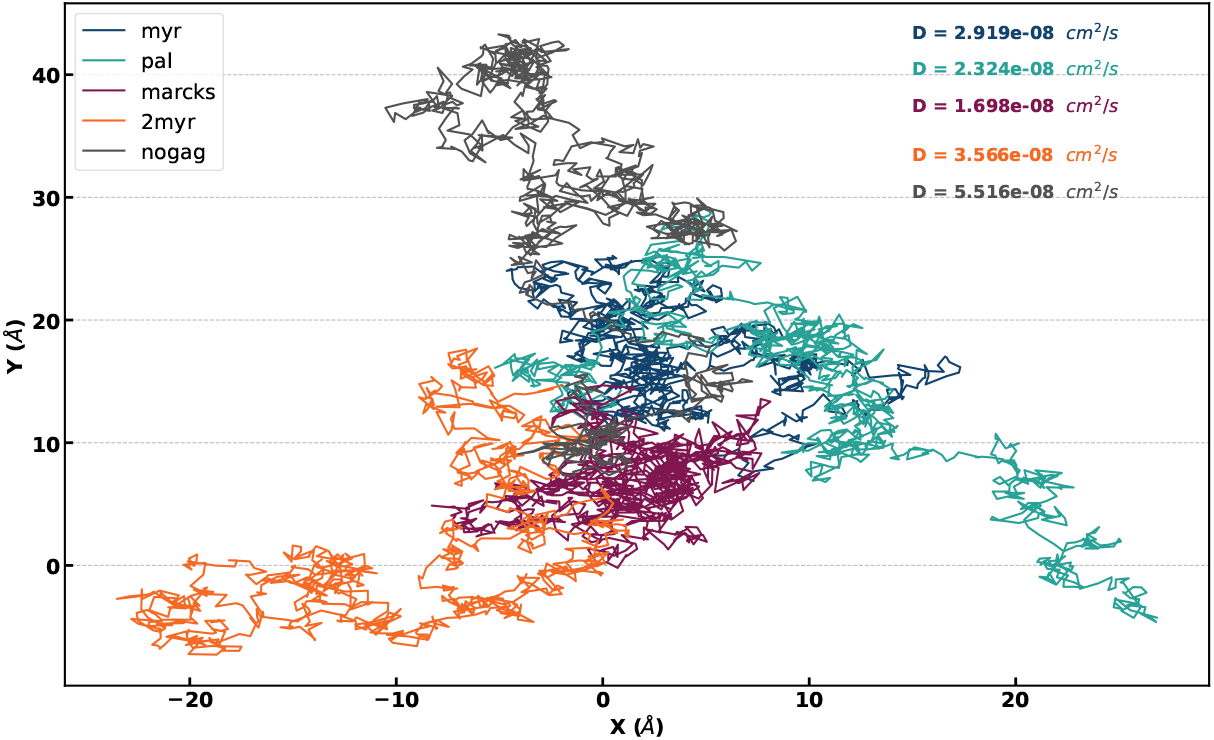
Trajectories of trimeric protein movements on a two-dimensional lipid membrane surface. The paths for different models are shown, including I (dark blue), II (teal), III (purple), IV (orange), and V (black). Each trajectory depicts the spatial exploration in the X-Y plane (in angstroms, Å) over time. The diffusion coefficients (D) for each trimer are listed, indicating their relative rates of lateral movement on the membrane. These coefficients help illustrate the variability in membrane mobility across different trimer types.

Further investigation, combined with the probability distribution function (PDF) of protein-membrane distance (Figure 15), reveals that lipid tail insertion also significantly decreases the distance between the protein platform and the membrane. This effect is especially pronounced in models with more than two lipid tails successfully integrated into the membrane. Specifically, in models I, II, and III, the distances are markedly reduced compared to models IV and V. According to Figure 15, the distances for the first three models are 47.17, 46.59, and 46.92 Å, respectively, whereas in models IV and V, the distances are 48.53 and 50.74 Å, respectively.

**Figure 15:**
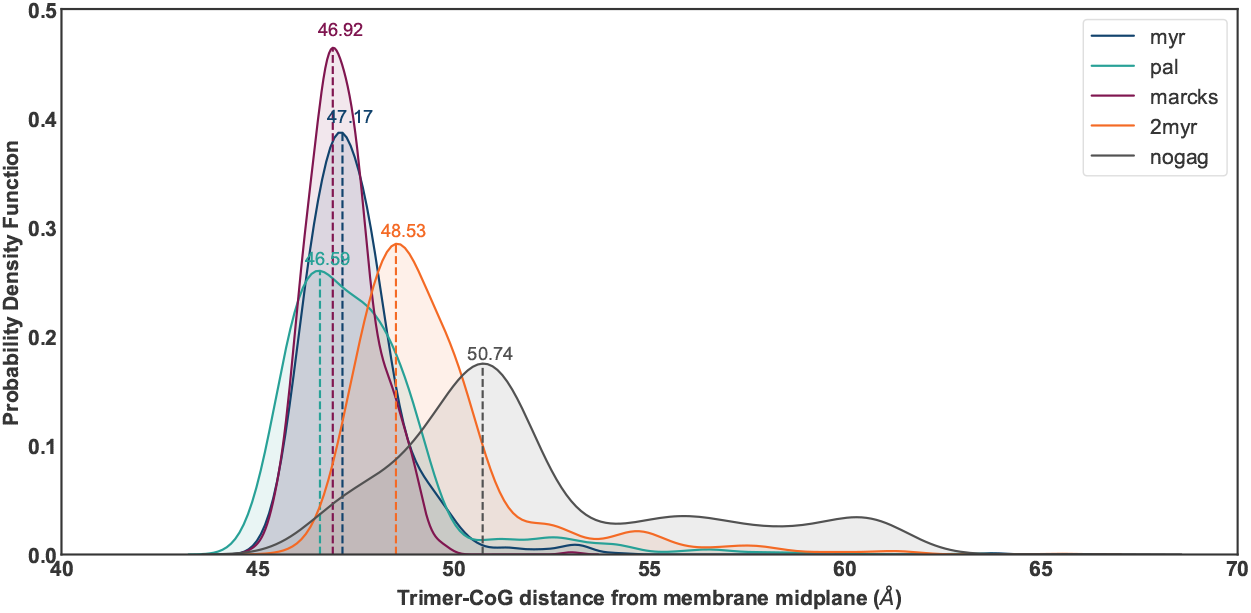
Probability distribution function showing the distance between the center of Geometry (CoG) of various trimer models and the membrane midplane (measured in angstroms, Å). The plot compares different models, including including I (dark blue), II (teal), III (purple), IV (orange), and V (black). The peak positions represent the most probable CoG distances for each model, with numerical labels indicating the specific distance values. This illustrates the differences in positioning among the various trimer models relative to the lipid membrane.

Taken together, these results highlight the critical role of lipid tails in modulating the interaction between the protein’s HBR and the charged lipids within the membrane. While electrostatic interactions bring the protein close to the membrane, the insertion of lipid tails reduces the protein’s lateral mobility, allowing it to remain in a more confined local lipid environment. This immobilization acts as another type of biophysical ‘cheap trick,’ reduction of dimensionality [31], and provides sufficient time for charged lipids to migrate from other regions of the membrane and accumulate at the protein’s HBR sites. As more lipids accumulate, electrostatic sequestration can occur, decreasing the average distance between the protein platform and the membrane and stabilizing the interaction between charged lipids and protein binding sites. This stabilization reinforces the protein-lipid complex and enhances the membrane curvature generation process.

Finally, it can be said that our results reveal that these two mechanisms operate synergistically in a feedback loop (Figure 13). The stronger the interaction at the highly basic region (HBR) increases the likelihood of lipid tail insertion into the membrane. Increased lipid tail insertion reduces protein mobility, which in turn amplifies electrostatic interactions by allowing more charged lipids to accumulate around the protein. Additionally, lipid tails further strengthen these interactions by decreasing the average distance between the protein platform and the membrane, thereby reinforcing the protein-lipid complex. This mutual reinforcement promotes greater membrane curvature. The optimal performance is observed in model III, which exhibits the lowest diffusion coefficient and shortest protein-membrane distance.

These results align with direct curvature measurements induced by this protein (Figure 5), underscoring the cooperative role of the HBR and lipid tails in driving membrane curvature.

The elucidated mechanism by which each triangular trimer induces curvature provides insight into how the entire nano-cage similarly imparts curvature to the membrane. Specifically, the arrangement of trimers in the nano-cage forms hollow pentagonal structures, with each trimer positioned at an angle of 58.3 degrees relative to its landing plane (Figure 1.A). This configuration places attraction points at the vertices of these pentagons. Since the rigidity of the nano-cage is higher than that of the membrane, it is assured that this mechanism effectively induces curvature in the membrane without causing any bending within the nano-cage itself.

The results from our models demonstrate that an increase in electrostatic interaction energy at the N-terminal significantly enhances the protein trimer’s efficiency in inducing membrane curvature. Which, at the experimental scale, translates to higher efficiency in the production of vesiculated nanocages. Notably, these findings are consistent with experimental data reported in the literature [50]. For instance, studies have shown that substituting the sequence of the HIV-1 Gag protein in membrane domain, which has a release efficiency of 13%, with those from the MARCKS protein reduces the vesicle release efficiency to 5%. Conversely, substituting these amino acids with the PH domain from PLC_*δ*_ enhances it to 15%.

Theoretical predictions align with these experimental results, suggesting that increasing the charge in the membrane-binding domain of the PLC_δ_-PH model to +4, compared to +2 in the HIV-1 Gag model, enhances release efficiency. In the MARCKS model, although its protein’s effector domain carries a high charge of +13 (as shown in Figure 16), it readily detaches from the membrane in the presence of Ca^++^/calmodulin or upon phosphorylation of three serine residues by PKC [31]. This detachment weakens its interaction with the membrane, subsequently reducing curvature induction. Similarly, in Model IV, the presence of a second tail disrupts the membrane interaction of the binding domain, leading to a decrease in curvature generation efficiency.

**Figure 16:**
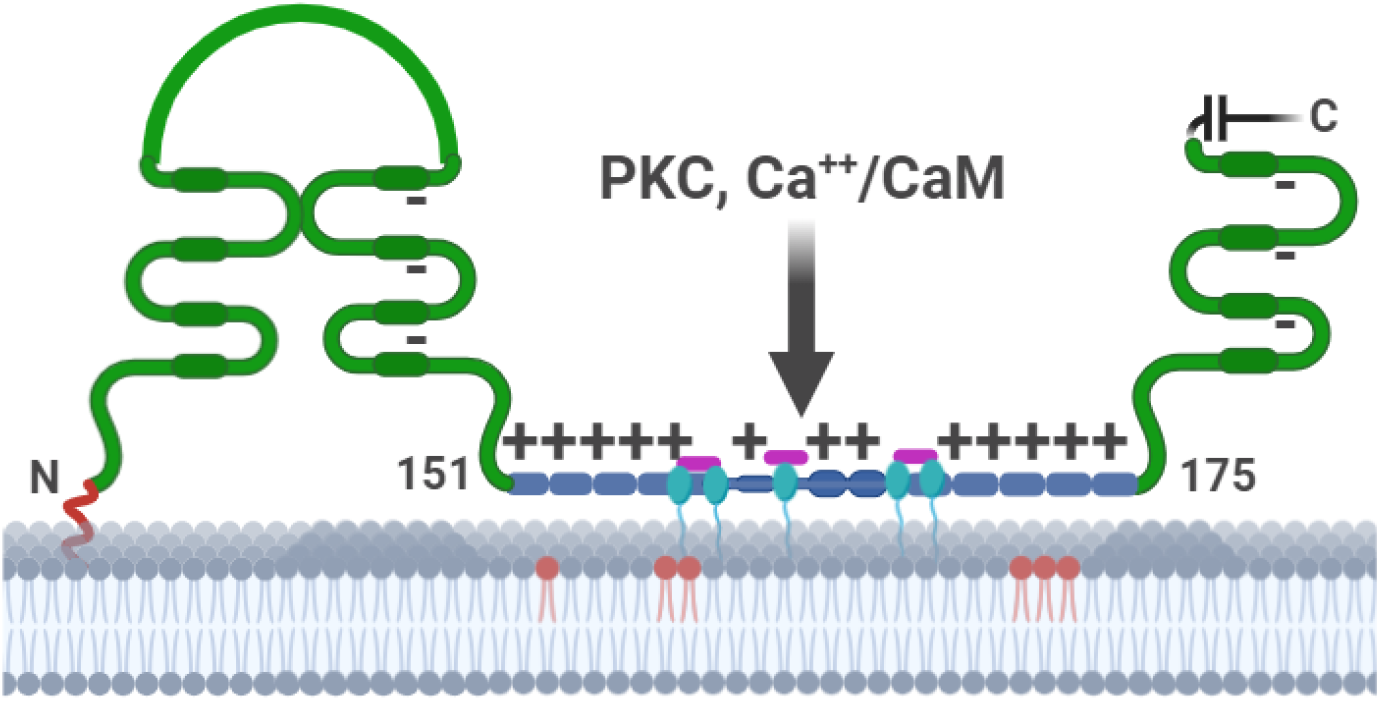
Illustration of MARCKS bound to a lipid bilayer. The myristoyl group (shown in red) integrates into the membrane through hydrophobic interactions, while the 13 basic residues (depicted in blue) in the effector domain interact with acidic lipids, represented by 6 red circles indicating PIP_2_. Additionally, five phenylalanine residues (depicted in cyan) are embedded in the bilayer. Phosphorylation of the three serine residues (highlighted in purple) by PKC or the binding of Ca^++^/calmodulin leads to the displacement and detachment of the effector domain from the bilayer. Inspired by [31]

Overall, The strong agreement between our simulations and experimental observations reinforces the validity of our findings and provides deeper insights into the biophysical mechanisms underlying these interactions.

With the mechanism of curvature induction now clarified, it remains essential to determine the specific contribution of each trimeric triangle’s interaction with the membrane to the total energy required for nano-cage vesiculation.

For this purpose, PMFs were further reconstructed from extensive MD simulations using Jarzynski’s method. The distance between the protein platform (excluding its non-structural N-terminal region) and the mid-plane of the membrane is used as the reaction coordinate (Figure 17).A plateau is observed in the PMF plot at a distance of approximately 65 Å, corresponding to an energy cost of around 280 kcal/mol (see Figure 17). At this point, the detachment of one lipid tail occurs concurrently with the separation of the protein core from the membrane.

**Figure 17:**
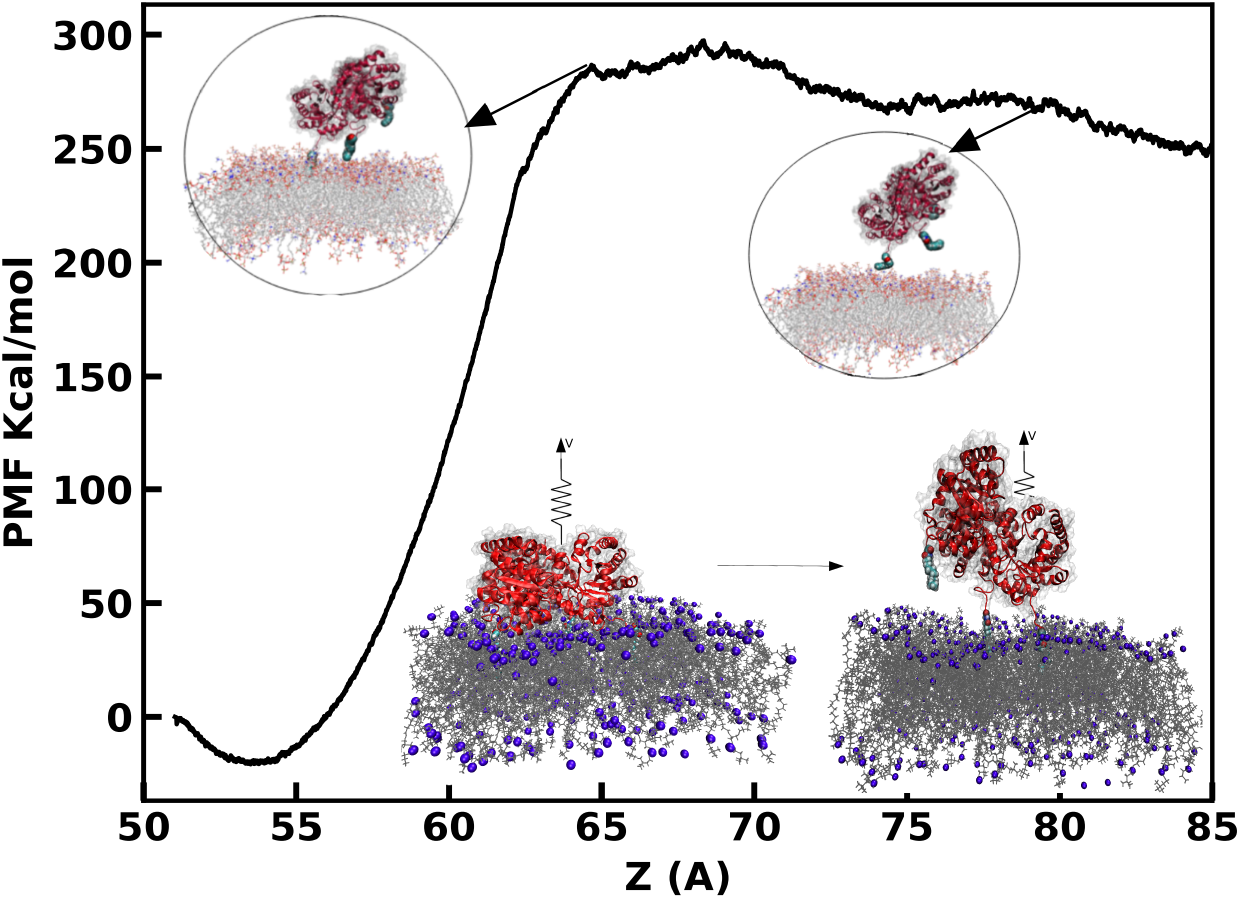
Potential of mean force (PMF) profile for the detachment of the protein from the membrane along the Z-axis. The PMF rises sharply as the protein core begins to separate from the membrane, reflecting the high energy required to overcome van der Waals and electrostatic interactions. The peak in the PMF corresponds to the detachment of the protein core and initial extraction of lipid tails. The insets illustrate key stages in the detachment process, showing shifts in protein position and membrane interaction. The gradual decline in PMF after the peak indicates the completion of lipid tail extraction and the protein’s release from membrane constraints.

**Figure 18:**
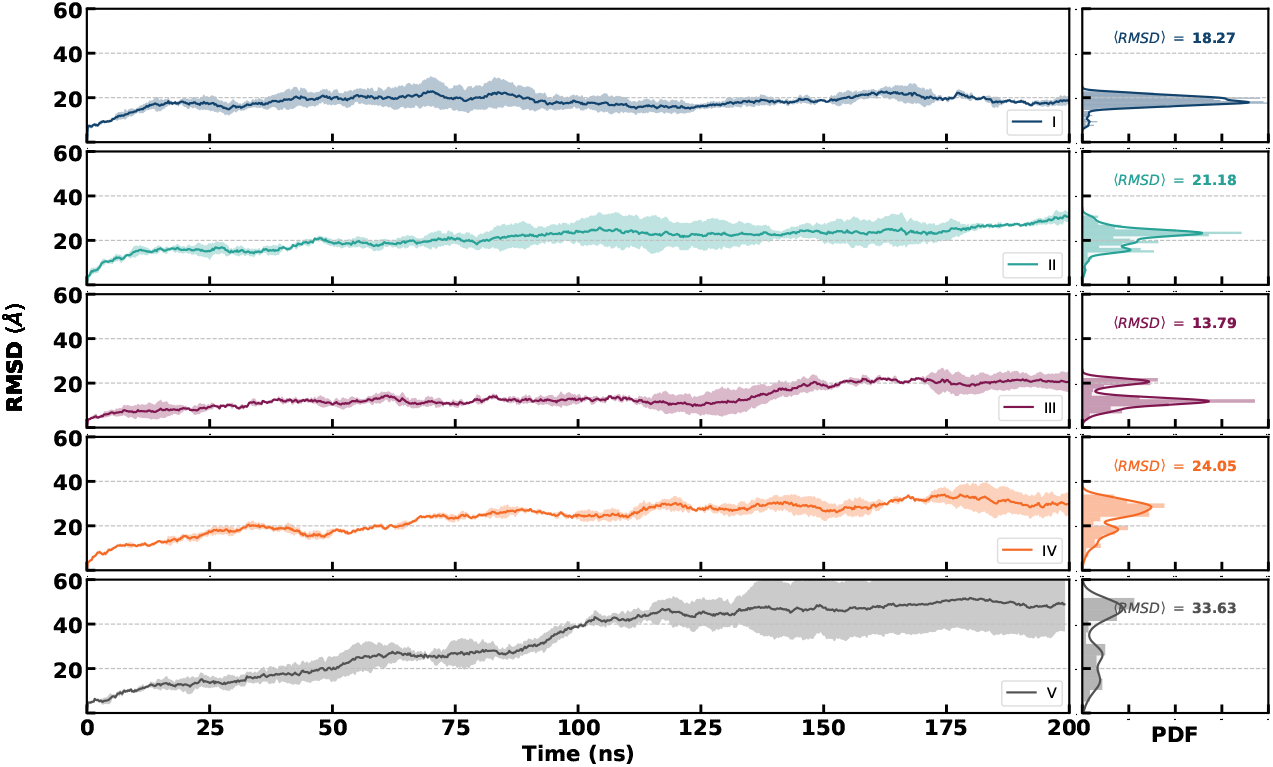
RMSD variation of the models during 200 ns of simulation. Each plot shows the root mean square deviation (RMSD) over time for different models (I to V), with the corresponding probability density function (PDF) of the RMSD values shown on the right. The average RMSD (⟨RMSD⟩) is indicated for each model, highlighting the structural stability and variations observed across the simulations. Models I and III exhibit clear plateaus after approximately 100 ns, indicating that these structures have reached stable configurations. Model II also shows relatively minor fluctuations after 100 ns, suggesting it is nearing equilibrium. For Models IV and V, while there is a continuous increase in RMSD earlier in the simulation, both display plateau regions after around 175 ns, indicating that they may be approaching a more stable configuration toward the end of the simulation.

Further progression of the process led to a decrease in the PMF, indicating that less energy is required to detach the remaining lipid tails compared to the protein core. At a distance of approximately 80 Å, all three lipid tails had fully dissociated from the lipid membrane. The energy data suggest that each trimer binding to the membrane contributes around 280 kcal/mol, facilitating membrane curvature.

This binding energy can be compared with the estimated energy required for vesiculation, calculated using the Helfrich Hamiltonian [11]:

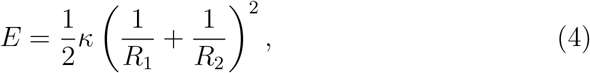

where *R*_1_ and *R*_2_ represent the local radii of curvature, and *κ* denotes the bending modulus of the membrane. For a typical phospholipid bilayer, *κ* is approximately 20 *k*_*B*_*T*, with *k*_*B*_*T* ≈ 4.1 × 10^−21^ J ≈ 0.6 kcal/mol [39].

Thus, the total energy required to form a spherical membrane vesicle of radius *R* can be estimated as:

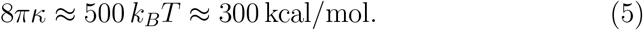

Comparing these values reveals that each trimer binding to the membrane, providing approximately 280 kcal/mol, contributes nearly sufficient energy for vesiculation. Since the energy required to induce vesiculation is around 300 kcal/mol, a single trimer binding is almost enough to achieve this transformation. Consequently, we infer that the binding of one or, at most, two trimers to the membrane should provide the necessary energy to drive vesiculation efficiently. However, experimental results not only support these estimations but also indicate that, on average, approximately 14 nanocages of type I are required to form a complete vesicle.

These experimental results indicate that an overly simplified application of the Helfrich Hamiltonian may not fully capture the complexity of this phenomenon. Numerous studies [11, 10] have shown that the simultaneous interaction of multiple proteins with the membrane can induce additional forces between proteins, beyond the direct interaction forces they experience with the membrane itself. These additional forces arise due to local curvature generated within the membrane, effectively transmitting disturbances to more distant points. In this way, lipid membranes act as a medium that propagates localized deformations, much like electric fields mediate interactions between charges or the curvature of space-time mediates gravitational interactions. However, unlike these fields, the membrane-mediated field in this context is tangible and directly observable.

It is important to note that this study has focused on the mechanism of curvature induction at the level of individual protein trimers. We have not yet addressed the potentially significant cooperative effects that may emerge from the simultaneous interaction of multiple nanocages with the membrane. Quantifying these cooperative effects, and determining their influence on the overall energy landscape of vesiculation, represents a crucial direction for future investigations, particularly for understanding the collective behavior of nanocages in membrane remodeling and vesicle formation.

## Conclusions

In recent years, synthetic protein nanocages have emerged as powerful tools in various fields, particularly in pharmaceuticals and medical applications. Inspired by the delivery mechanisms of viruses like HIV-1 and hepatitis, which efficiently transport genetic material to host cells, researchers have sought to replicate these strategies, transforming synthetic nanocages into functional artificial viruses. Our study provides insight into the molecular mechanisms underlying these transformations, offering a foundational understanding necessary for the optimal design of hybrid biological materials.

The first step toward creating robust and effective nanocages is a deep understanding of the molecular processes that lead to vesiculation. Through molecular dynamics simulations, we investigated these processes, focusing on how certain lipid and protein configurations promote membrane curvature. Our results reveal that membrane curvature is primarily driven by the accumulation of longer lipids at the peripheral regions of the protein platform, leaving the central areas devoid of these lipids. This lipid redistribution creates tension and curvature across the membrane, which is a critical factor in forming vesicular structures.

Among the models studied, those with stronger highly basic regions (HBR) and specific lipid-binding sites showed the greatest ability to induce membrane curvature. In these models, the interaction between the HBR and the membrane facilitates the insertion of lipid tails into the membrane, anchoring the protein to the lipid bilayer. This, in turn, reduces protein diffusion, allowing a higher concentration of negatively charged lipids to gather around the protein. Such configurations result in a shorter distance between the protein platform and the membrane, enhancing the stability and strength of the protein-lipid complex (model I to model III).

Our findings also show that any interference in lipid tail insertion, even when the HBR is structurally similar to other models, significantly reduces the protein’s ability to induce membrane curvature. This decline in function underscores the importance of a well-coordinated interaction between the HBR and lipid tails for achieving optimal curvature (*C*_*IV*_ < *C*_*I*_). These results suggest that the HBR and the lipid-binding sites work synergistically, reinforcing each other’s roles in a stepwise, feedback-controlled loop. This mutual reinforcement plays a critical role in enabling the protein complex to induce membrane curvature effectively, as illustrated in Figure 13.

The alignment of our simulation results with experimental data further validates our analysis, providing a reliable model for studying the molecular mechanisms involved in vesiculation. By measuring the binding energy of each protein trimer with the membrane, we quantified the contributions of each component of the nanocage to the energy required for vesiculation. We estimated a binding energy of approximately 280 kcal/mol per trimer, which, in theory, exceeds the energy necessary for vesiculation and the formation of a spherical lipid coating around the nanocage.

In real biological conditions, where many proteins are present on the membrane surface, additional factors such as entropic forces from the membrane, alongside other presenting and known nanoscale forces, come into play, influencing the curvature formation process. Although these elements extend beyond the scope of the current study, they are essential for a complete understanding of membrane dynamics and will be addressed in future research. In subsequent work, we will explore how these forces impact protein packaging and the stability of vesicular structures, ultimately advancing our knowledge in the design of synthetic nanocages for biomedical applications.

## Appendix

To differentiate the curvature induced by the protein trimer from thermally-induced random deformations of the bilayer surface, we calculated the local curvature of the bilayer over time. The membrane surface was initially modeled by fitting the coordinates of the phosphorus beads of the lipids to a two-dimensional fifth-degree polynomial:

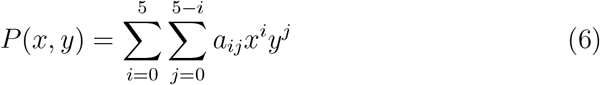

Here, the outer sum iterates over the power *i* of *x* from 0 to 5, and the inner sum iterates over the power *j* of *y* from 0 to 5 − *i*, ensuring that the total degree *i* + *j* does not exceed 5. The coefficients *a*_*ij*_ correspond to each term *x*^*i*^*y*^*j*^.

Since the surface lacks steep gradients and the induced curvature of the bilayer is minor (Figure 6.B), the mean curvature *C*(*x, y*), can be linearly approximated using the Laplacian of the height function, as described by the Monge equation:

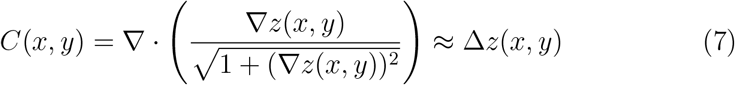

The local curvature per unit area beneath the protein trimer is then defined as:

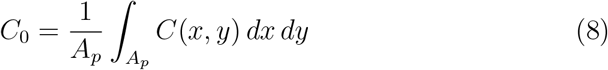

where *A*_*p*_ is the projected area of the trimer on the membrane surface. In this formulation, inward bending is represented by a positive *C*_0_, while outward bending is indicated by a negative *C*_0_.

## Author contributions

S.R. conducted the simulations and data analysis, and prepared all plots and figures for the manuscript. S.R. and M.R.E. conceived and designed the research project, and jointly contributed to the interpretation of the results.

S.R. drafted the main manuscript text. All authors critically reviewed and approved the final manuscript.

## Conflicts of interest

Conflicts of Interest: The authors declare that they have no conflicts of interest to disclose.

## Data availability

The molecular dynamics simulation trajectories are available from the corresponding author upon reasonable request. Due to the large size of the raw trajectory files, direct public deposition is not feasible, but processed data and analysis scripts are provided in the Supplementary Information.

## Acknowledgements

We are grateful to the HPC of SUT to provide the computational resources of this study.

## Notes

### Competing Interest Statement

The authors have declared no competing interest.

